# Behavioral signatures suggest distinct modes of suppressing irrelevant information during tactile temporal attention in human participants

**DOI:** 10.64898/2026.08.29.747844

**Authors:** M. Gironimi, M.E. Diamond

**Affiliations:** SENSEx Laboratory, International School for Advanced Studies (SISSA), 34136 Trieste, Italy

## Abstract

To make adaptive perceptual judgments, the nervous system must selectively process behaviorally relevant sensory information while filtering out competing distractions. Although attentional control has been extensively studied in the visual domain, particularly in the context of spatial selection, considerably less is known about how attention operates in the tactile modality and across time rather than space. Here, we developed a paradigm to investigate temporal tactile attention in human participants, enabling the study of attentional behavior and underlying behavioral strategies in this sensory domain.

Participants were instructed to categorize the intensity of a task-relevant tactile stimulus delivered to the fingertip while ignoring an irrelevant tactile stimulus. A visual cue indicated which of two sequentially presented stimuli was relevant on each trial. In addition, participants completed self-report questionnaires assessing autistic traits and aberrant salience (the tendency to assign significance to otherwise neutral stimuli or events).

Across subjects, participants performed the task with high accuracy. However, clustering analyses based on behavioral features revealed distinct response profiles. One cluster did not exhibit biases induced by the irrelevant stimulus. In contrast, a third cluster displayed a repulsive effect of the irrelevant stimulus and showed lower overall performance. These behavioral phenotypes were also reflected, to some extent, in differences in learning trajectories across training sessions.

We further explored participants’ metacognitive awareness through a post-experiment questionnaire assessing subjective evaluations of task difficulty and performance. Although exploratory, the results suggest a relationship between metacognitive reports, behavioral strategies, and objective task performance. In contrast, neither autistic traits nor aberrant salience scores were associated with performance measures or behavioral phenotypes.

Together, these findings support the view that attentional control is implemented through multiple, individualistic behavioral strategies rather than a single mechanism pertaining to all, suggesting that task structure interacts with individual predispositions to shape distinct modes of attentional control in human participants.

## Introduction

Goal-directed behavior depends on a set of higher-order cognitive command processes collectively referred to as executive functions. Among these, attentional control enables organisms to prioritize behaviorally relevant information while suppressing competing or distracting inputs. By regulating access to limited processing resources, attention plays a fundamental role in perception, decision-making, and adaptive behavior.

Over the past decades, attentional control has been extensively investigated from both cognitive and neuroscientific perspectives. Research on attention has been largely dominated by the visual modality and by paradigms examining the allocation of attention across space [1–3]. Classical theories focused on the locus and mechanisms of attentional selection, proposing models of early selection, late selection, and attentional attenuation [4–6]. Subsequent neurophysiological and neuroimaging studies identified the role of top-down modulation from the frontoparietal attentional networks to early and extrastriate visual cortices [7–9].

Considering attention only in the context of vision and spatial selection is a narrow perspective. Organisms must often prioritize information that unfolds over time and across multiple sensory modalities. Touch represents a particularly relevant domain. Beyond its role in object exploration and manipulation [10], tactile processing is essential for social interactions [11], and becomes especially important under conditions of sensory deprivation, such as blindness, where it can support complex perceptual functions including Braille reading [12, 13]. Nevertheless, the mechanisms governing attentional control in the tactile modality remain considerably less explored than those described for vision.

One of our previous studies [14] showed that rodents implement multiple behavioral and neuronal strategies when performing a tactile temporal attentional task, that emerged under the same task demands. However, it is still unclear whether attentional control in human participants is expressed through alternative solutions shaped by both task structure and individual predispositions.

With the present study, we sought to establish whether human participants implemented attentional control through a common mechanism, or whether multiple modes of suppressing irrelevant information would emerge. More specifically, we asked whether variability among participants would be primarily quantitative, reflecting differences in overall performance, or whether qualitatively distinct modes of interaction with irrelevant information would also be observed.

Participants performed a task in which two sequential tactile stimuli were presented and a visual cue indicated which stimulus was relevant for the perceptual judgment. Participants were asked to categorize the intensity (weak or strong) of the relevant stimulus. Four blocks of trials were presented, distributed over two experimental sessions. This design allowed us to characterize not only steady-state performance but also learning trajectories over time, facilitating direct comparisons with the behavioral patterns previously observed in rats.

In addition to the behavioral task, participants completed self-report measures assessing autistic traits and aberrant salience. Atypical attentional and executive processes have frequently been reported in autism spectrum disorder, while aberrant salience, defined as the inappropriate assignment of significance to otherwise neutral stimuli, has been linked to altered perceptual and cognitive experiences [15, 16]. However, whether variation in these traits contributes to attentional performance in non-clinical populations remains uncertain [17]. We therefore explored whether inter-individual differences in these dimensions were associated with task performance or with specific behavioral strategies.

Finally, we examined participants’ metacognitive awareness through a post-experimental questionnaire assessing subjective evaluations of task difficulty and performance. Although exploratory, this analysis allowed us to investigate whether differences in objective behavior were accompanied by corresponding differences in self-evaluation and introspective accuracy.

Overall, our aim was not only to characterize human performance in a temporal tactile attention task, but also to determine whether distinct attentional strategies emerge across individuals. Demonstrating such variability would provide evidence that attentional control is reflected through meaningful alternative behavioral solutions, rather than mere noise around a common mechanism.

## Results

### Temporal attention in the tactile modality of humans

To assess how humans implement tactile temporal attention, we tested 30 participants in a task requiring them to selectively attend to a relevant vibrotactile stimulus (REL) while ignoring an irrelevant stimulus (IRR), allowing us to investigate how attention is flexibly allocated across time in the human tactile modality.

Participants received vibrotactile stimuli on the right index finger, and were required to judge whether the relevant stimulus was "strong" or "weak", reporting their perceptual judgment by pressing one of the two response buttons (Figure 1a-b). Visual feedback was provided after every trial.

**Figure 1:**
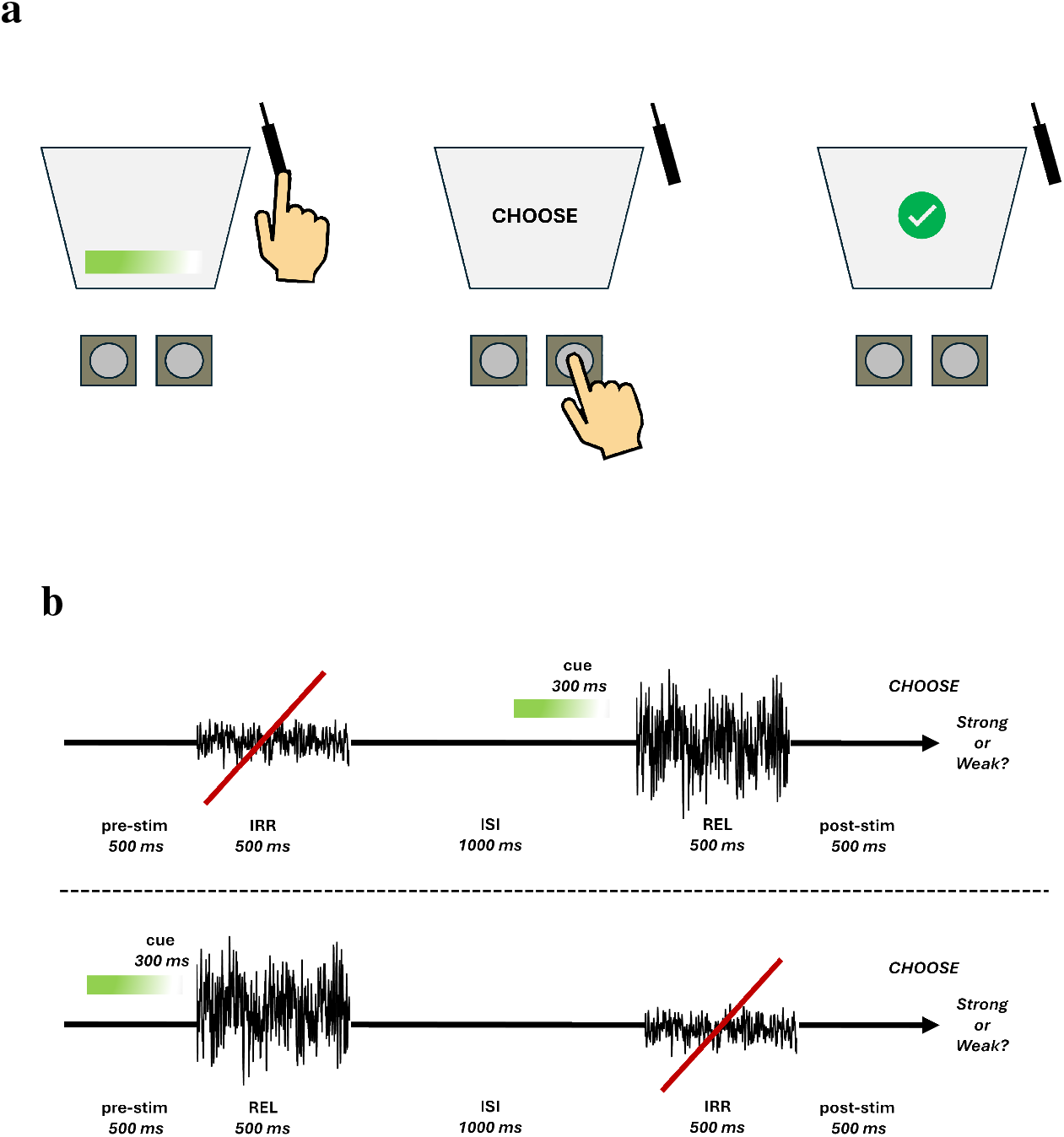
Setup configuration and task schematization. **a.** Experimental setup used for human testing. Vibrotactile stimuli were delivered to the participant’s right index fingertip while a visual cue identified the relevant stimulus. Participants categorized stimulus intensity by pressing one of two response buttons and received visual feedback after each trial. **b.** Structure of the task: the auditory relevance cue used in rats was replaced by a visual cue, and the inter-stimulus interval was fixed at 1000 ms.

We used a stimulus set consisting of seven relevant-stimulus intensities and five irrelevant-stimulus intensities. Each vibration consisted of a sequence of velocity values sampled from a Gaussian distribution and was defined by its nominal speed (mm/s).

Before testing, participants completed a demographic questionnaire reporting age, gender, nationality, neuropsychiatric diagnoses, and current pharmacological treatment. Following data collection, participants reporting neuropsychiatric diagnoses or psychotropic medication use would have been excluded from the study; however, no exclusions were necessary. Participant anonymity was maintained through the use of alphanumeric identifiers.

Participants also completed three self-report questionnaires: the AQ-10, the SPQ-10, and the Aberrant Salience Inventory (ASI). These measures were collected to evaluate whether variability in autistic traits and aberrant salience was associated with performance in a task requiring flexible allocation of attention across time. Previous findings linked ASD (i.e., autistic spectrum disorder) traits and aberrant salience to attentional deficits [15], [18].

Testing was conducted across two experimental sessions separated by no more than one week. Each session included two testing blocks, for a total of four blocks per participant. Before the first session, participants completed a brief warm-up phase designed to familiarize them with the task and experimental setup. During this phase, they also performed 20 reference-memory trials, as in [19], to ensure they understood that stimulus categorization required comparison with a reference intensity. Participants who did not demonstrate adequate understanding of the task were allowed to repeat the warm-up phase once before continuing.

The overall structure of the study is illustrated in Figure 2.

**Figure 2:**
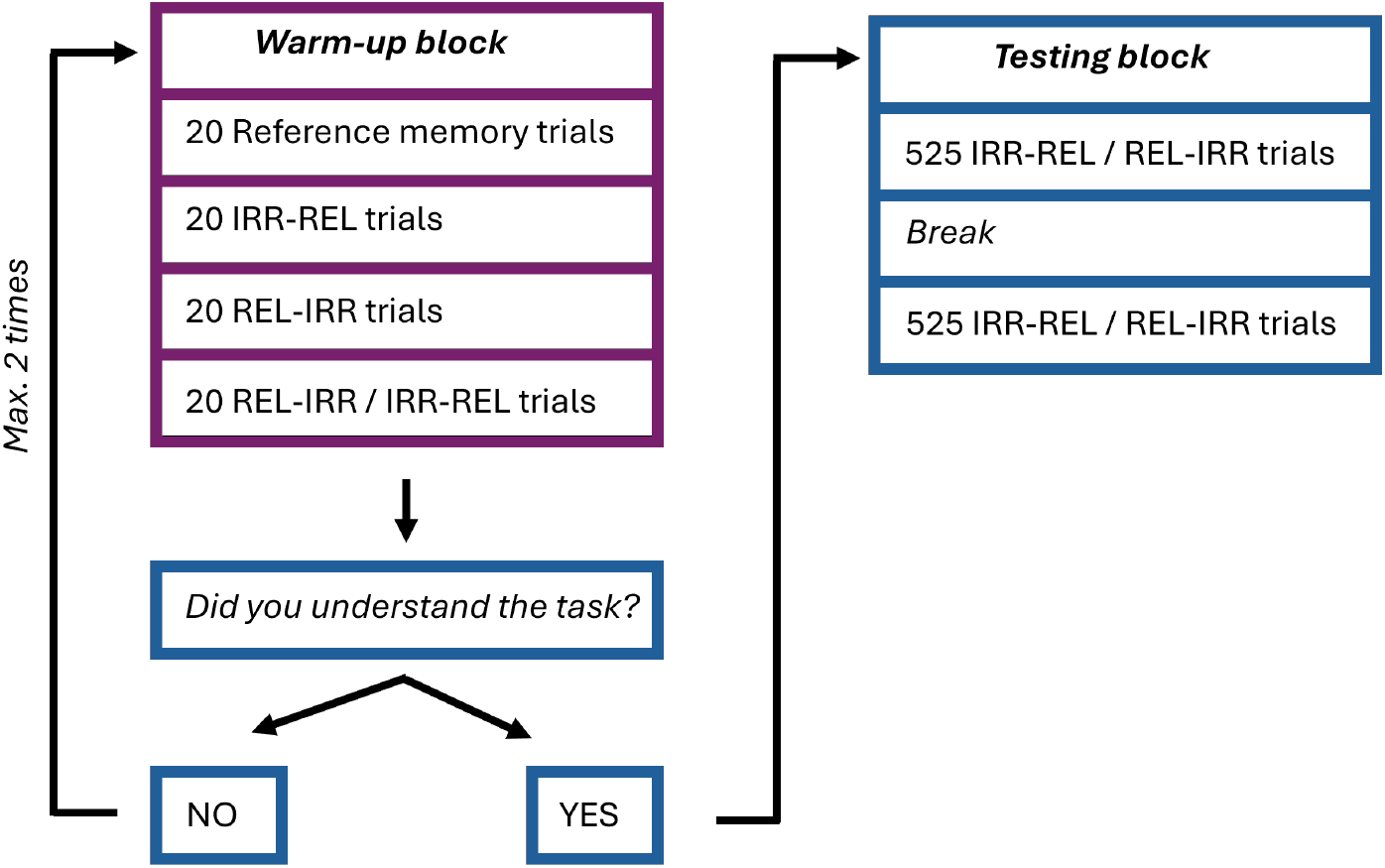
Structure of the human study. Participants first completed a warm-up phase that could be repeated once if necessary. After confirming task comprehension, they completed the testing blocks across two experimental sessions held on separate days. At the end of the second session, participants completed a brief post-task questionnaire designed to assess subjective perceptions of task difficulty and confidence. The questionnaire contained nine items with three response options each and evaluated perceived task difficulty, confidence in performance, the influence of stimulus position, and participants’ memory of the reference stimulus.

### Effect of irrelevant information on humans’ perceptual judgments

Participants showed high levels of performance in both trial types and were substantially less influenced by irrelevant information than were rats. Psychometric functions revealed accurate perceptual judgments during both REL-IRR and IRR-REL trials, indicating that participants successfully used the relevant stimulus to guide their decisions independently of order (Figure 3).

**Figure 3:**
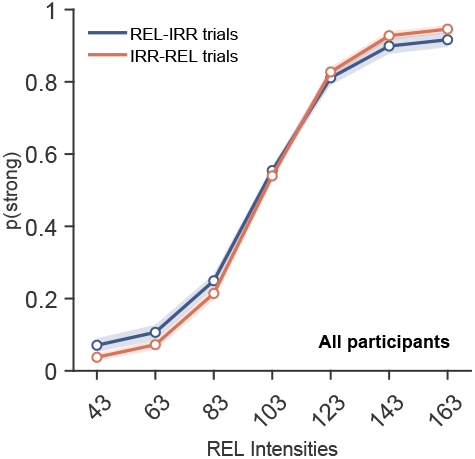
Across-subject overall performance. Psychometric functions showing average performance across participants for REL-IRR and IRR-REL trials. Dots represent the across-subject mean probabilities obtained from individually fitted psychometric functions at each tested REL intensity. Lines connect consecutive fitted probabilities for visualization, and shaded areas represent the S.E.M. across individually fitted probabilities. Corresponding empirical probabilities are shown in Supplementary Material.

Despite this overall high performance, conditioning behavioral responses on irrelevant-stimulus intensity revealed subtle effects of irrelevant information on perceptual judgments (Figure 4A). During REL-IRR trials, psychometric functions showed only minor differences across irrelevant-stimulus intensities, suggesting that participants largely ignored the irrelevant stimulus when the relevant stimulus appeared first. In contrast, during IRR-REL trials, the highest irrelevant-stimulus intensities produced a slight separation of the psychometric functions, whereby stronger relevant stimuli were more likely to be judged as weaker than their physical intensity would predict. This effect was largely absent during REL-IRR trials, consistent with a repulsive influence of the irrelevant stimulus.

**Figure 4:**
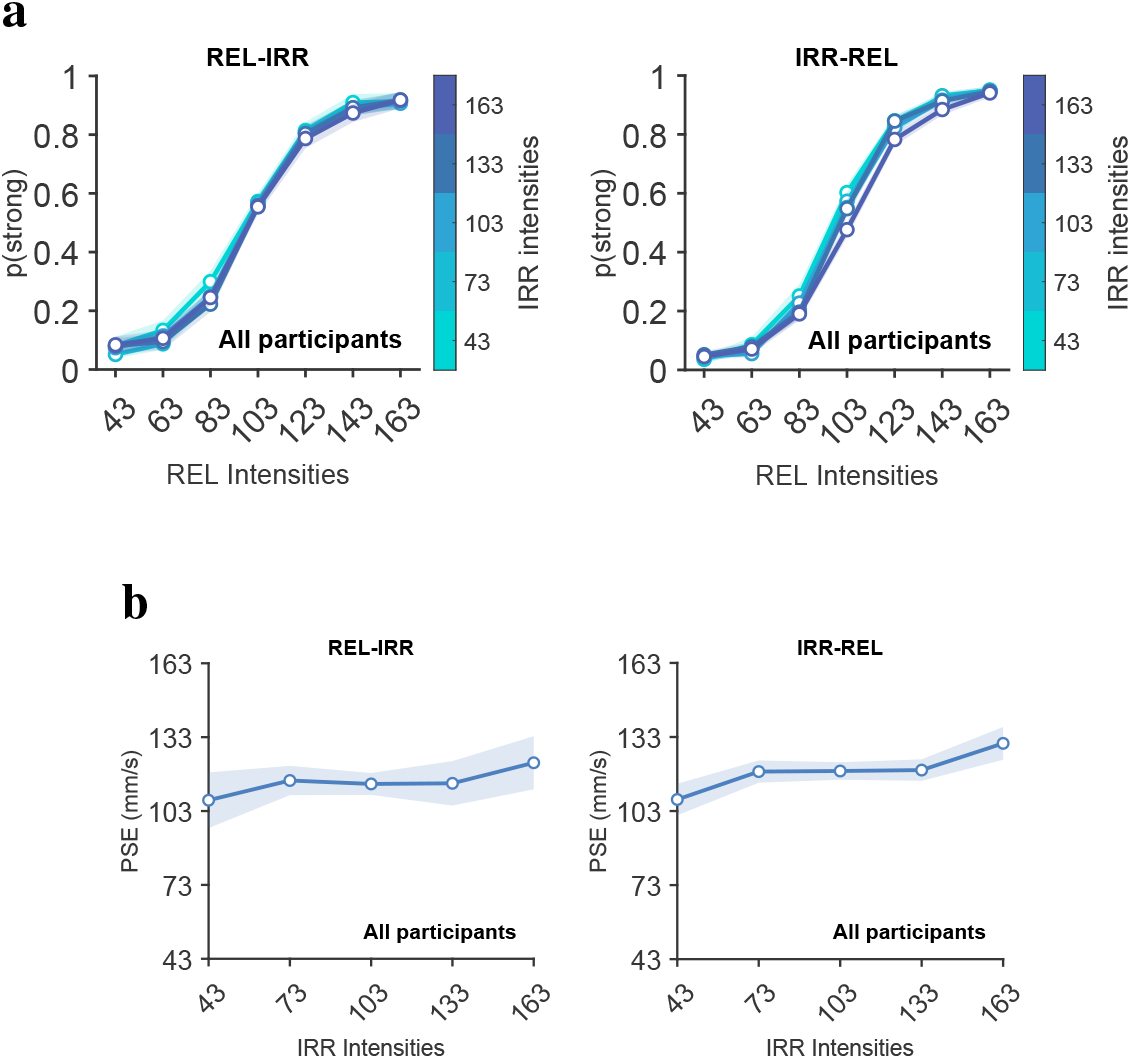
Across-subject irrelevant-stimulus bias. **a.** Psychometric functions conditioned on irrelevant-stimulus intensity during REL-IRR trials (left) and IRR-REL trials (right). **b.** Point of Subjective Equality as a function of irrelevant-stimulus intensity during REL-IRR trials (left) and IRR-REL trials (right). Shaded regions represent SEM of fitted PSE estimates.

To quantify these effects, we examined changes in the Point of Subjective Equality (PSE) as a function of irrelevant-stimulus intensity. PSE analysis revealed a positive relationship between irrelevant-stimulus intensity and PSE during IRR-REL trials (Figure 4b). As irrelevant stimuli became stronger, participants required increasingly stronger relevant stimuli to perceive them as equivalent to the reference stimulus.

This shift in PSE was consistent with the repulsive bias observed in the psychometric functions. Strong irrelevant stimuli altered perceptual judgments even when participants were instructed to ignore them, producing systematic changes in the perceived relationship between the relevant stimulus and the memorized reference value.

Overall, the effect of irrelevant information was modest across subjects. Participants generally performed the task accurately and successfully discounted distracting sensory input. Nevertheless, the presence of systematic PSE shifts and small but detectable psychometric biases indicates that irrelevant information continued to influence perceptual decisions under specific conditions, particularly when the irrelevant stimulus preceded the relevant one.

#### Changes in performance across the sessions

To assess whether participants progressively increased their reliance on the relevant stimulus (REL) and reduced the influence of the irrelevant stimulus (IRR), we examined the evolution of stimulus weights across the four testing blocks using generalized linear models. Across both trial sequences, the weight assigned to the relevant stimulus increased modestly over the course of training (Figure 5). This was observed in both REL-IRR and IRR-REL trials, indicating progressively greater reliance on task-relevant sensory information when making perceptual judgments.

**Figure 5:**
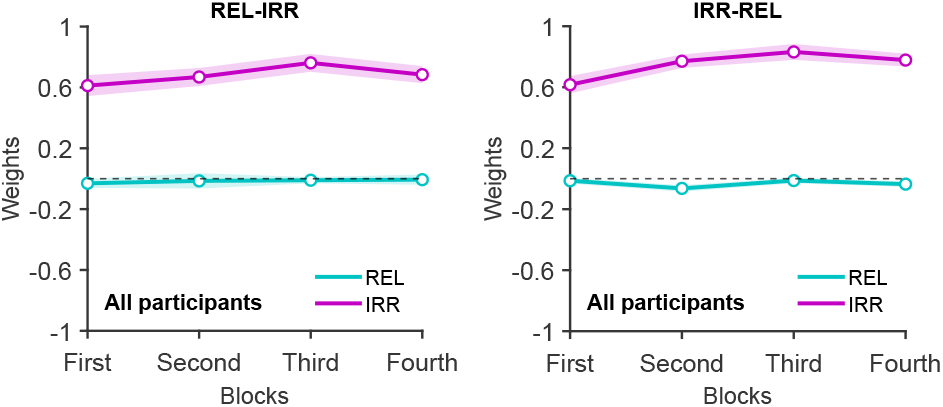
Across-subject learning progression. Generalized linear model weights for relevant (REL) and irrelevant (IRR) stimuli across the four testing blocks, shown separately for REL-IRR trials (left) and IRR-REL trials (right). Shaded regions represent SEM of fitted GLM weights.

In contrast, the weight assigned to the irrelevant stimulus remained close to zero across all four blocks (Figure 5). This pattern was consistent across trial types and indicates that participants successfully suppressed the influence of irrelevant sensory information throughout the experiment, in contrast to the rats.

Taken together, these findings suggest that participants mastered the task rapidly and relied primarily on the REL stimulus from the earliest stages of testing. The increased weight given to the REL stimulus may correspond to a very small increase in acuity – that is, perceptual plasticity.

#### Effect of previous-trial choices on perceptual judgments

Under some conditions, past choices exert an attractive influence on current decisions [20]. If such a bias occurred in our experiments, participants would be more likely to select “strong” on the current trial if they selected “strong” on the preceding trial, and vice versa. The data indicate that previous-trial choices exerted only a weak influence on current perceptual judgments when averaged across subjects.

Psychometric functions conditioned on the choice made in trial N-1 showed only a negligible attractive bias in both REL-IRR and IRR-REL trials (Figure 6). These results indicate that, at the population level, current perceptual judgments were driven primarily by information presented within the ongoing trial rather than by the choice made on the preceding trial.

**Figure 6:**
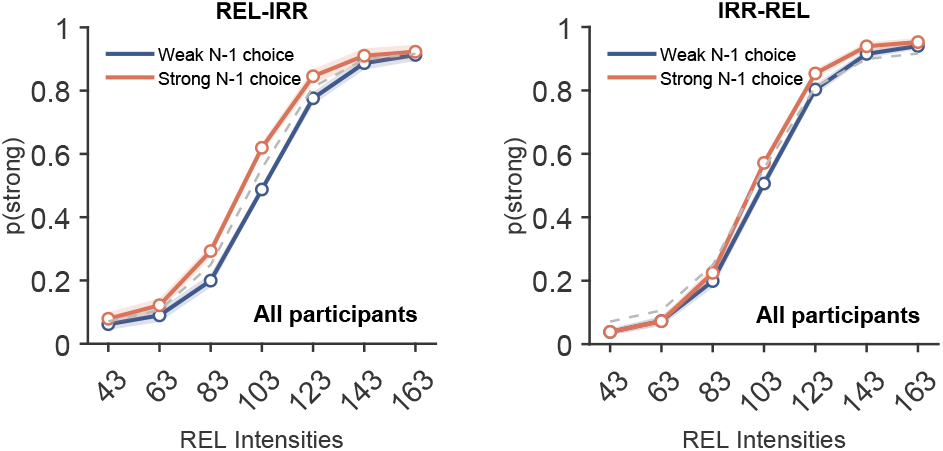
Effect of previous-trial choice on current perceptual judgments. Psychometric functions conditioned on the choice made in trial N-1, shown separately for REL-IRR trials (left) and IRR-REL trials (right).

### Behavioral clustering reveals distinct attentional strategies

#### Behavioral performance differs across clusters

When averaged across subjects, we identified a slight repulsive effect of IRR and a slight N-1 choice attractive bias. Because these tendencies both reflect the incorporation of irrelevant information, we posited that they may be somehow connected in individuals. We used these two behavioral metrics as clustering features, in order to divide the participants in three sub-pools. We performed a *k-medoids* clustering analysis, using *k=3*, to identify three clusters based on the GLM weights of trial N-1 choice and trial N IRR stimulus. Given the relatively small sample (n=30), we compared solutions using *k=2*, *k=3*, and *k=4*. The three-cluster solution provided a parsimonious separation into qualitatively distinct GLM weights profiles. Increasing to 4 subdivided one of these profiles into groups differing in weight magnitude, without revealing an additional qualitatively distinct pattern.

Specifically, for each subject, we computed two separate GLMs, calculating the weight of trial N-1 choice on trial N choice, and the weight of trial N IRR stimulus on trial N choice. These weights were then used to cluster participants in three groups. In Figure 7, Cluster 3 exhibited a strong IRR repulsive bias, with a repulsive trial N-1 choice effect. Cluster 2 showed a greater trial N-1 choice attractive bias, with a slight, overall repulsive effect of IRR. Cluster 1 exhibited less trial N-1 choice bias, and mixed attractive/repulsive IRR bias.

**Figure 7:**
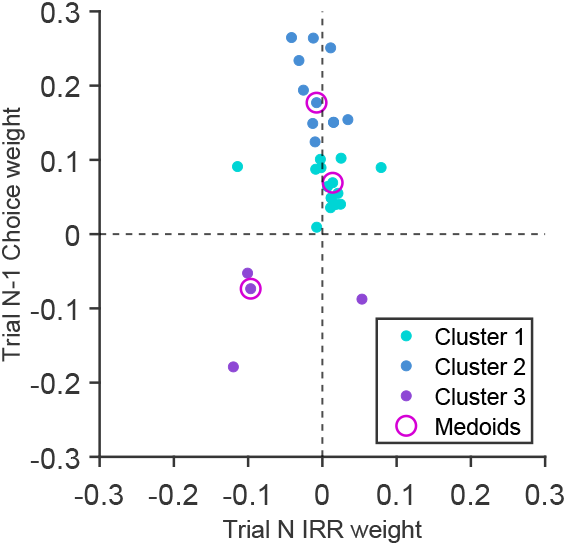
*k-medoids* clustering based on trial N-1 choice weight and trial N IRR weight. *k-medoids* clustering identified three distinct clusters of participants, with specific directions of trial N-1 choice and IRR bias.

Clustering revealed substantial differences in task performance and in the influence of irrelevant information on perceptual judgments. Although all groups successfully performed the task, the three clusters exhibited distinct behavioral patterns that were not apparent at the population level.

#### Clusters differ in performance and susceptibility to irrelevant information

Average task performance differed across clusters (Figure 8). Cluster 1 showed comparable performance in REL-IRR and IRR-REL trials, whereas Cluster 2 performed slightly better when the relevant stimulus appeared first (REL-IRR trials). Cluster 3 exhibited the lowest overall performance and, similar to the pattern observed in rats, showed reduced accuracy during REL-IRR trials relative to IRR-REL trials. These findings indicate that participants did not differ solely in the magnitude of their performance, but also in the trial types that posed the greatest difficulty.

**Figure 8:**
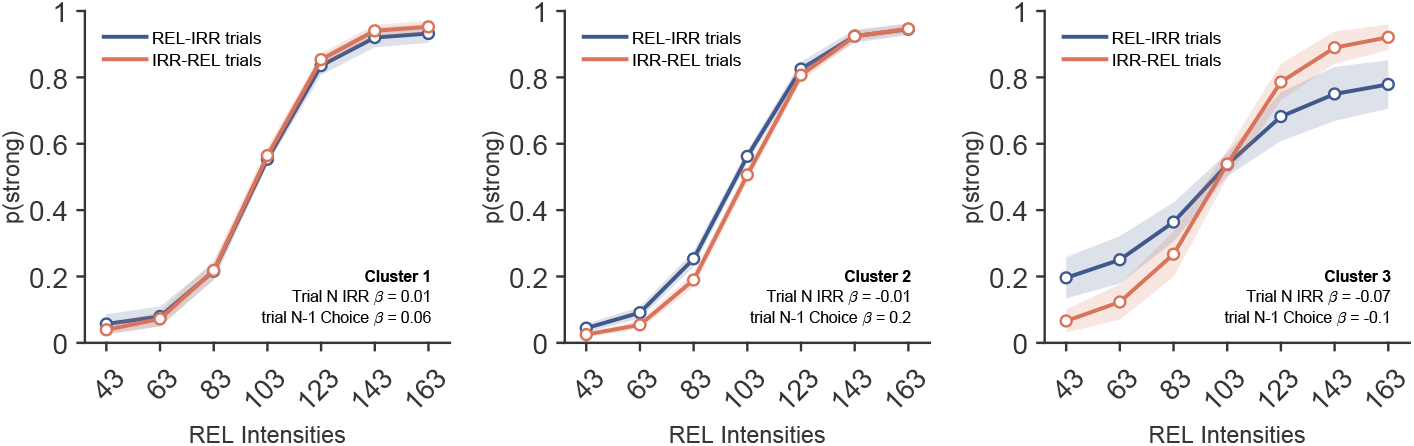
Average performance at the cluster level. Psychometric functions showing average task performance for Clusters 1, 2, and 3.

The clusters also differed in the effect exerted by the IRR stimulus (Figure 9). Cluster 1 showed psychometric functions that largely overlapped during REL-IRR trials and IRR-REL trials, indicating minimal influence of the irrelevant stimulus. This profile closely resembled the pattern observed in expert rats.

**Figure 9:**
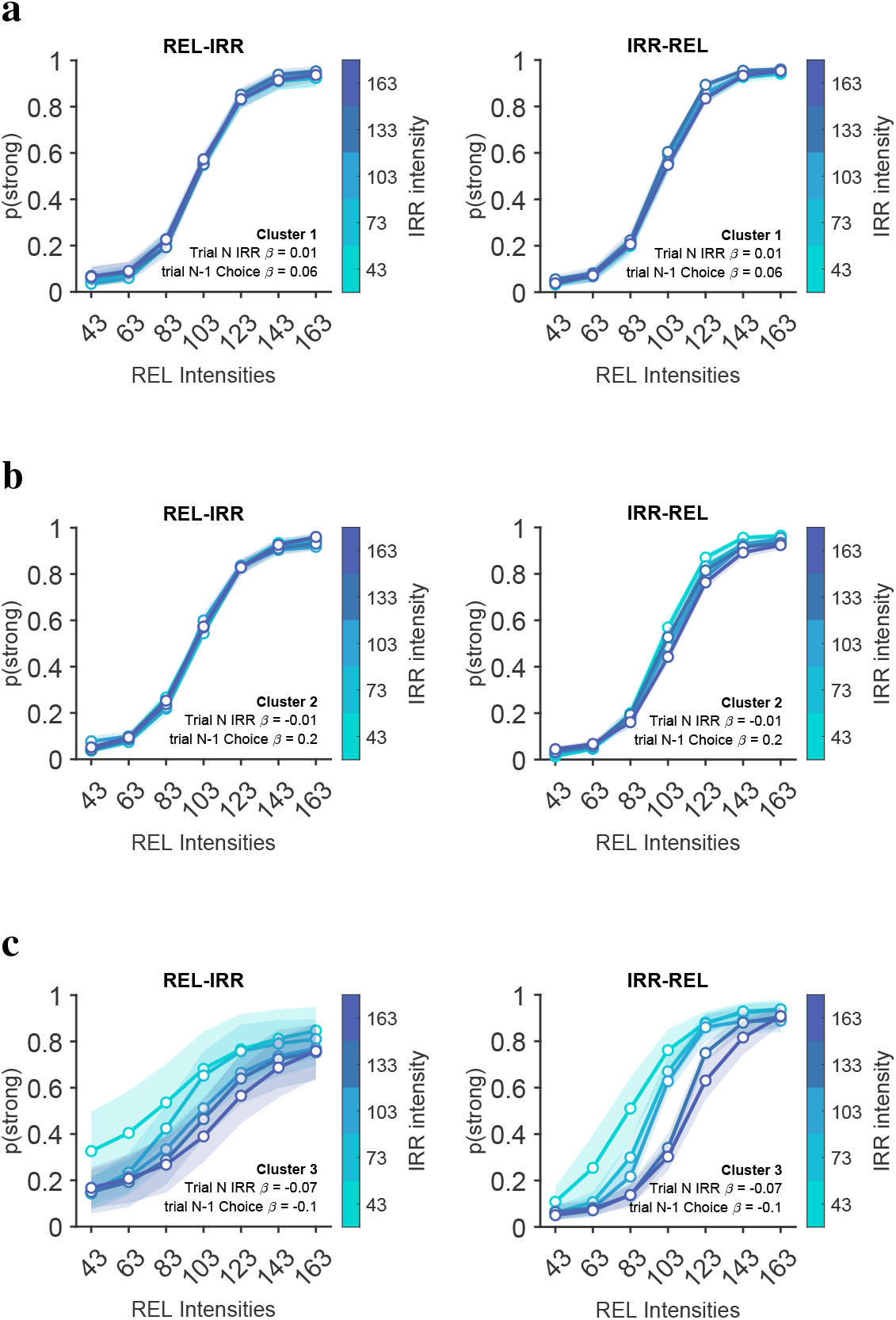
Irrelevant-stimulus bias at the cluster level. **a.** Psychometric functions conditioned on irrelevant-stimulus intensity for Cluster 1. **b.** Same analysis for Cluster 2. **c.** Same analysis for Cluster 3.

For Cluster 2, performance remained high across both trial types, but psychometric functions revealed a slight repulsive bias during IRR-REL trials. Irrelevant information tended to shift judgments away from, rather than toward, the intensity of the irrelevant stimulus.

Cluster 3 displayed a clear divergence from Clusters 1-2. Although participants in this cluster performed better during IRR-REL than REL-IRR trials, they exhibited a pronounced repulsive bias in both trial types, particularly when the relevant stimulus appeared first. This cluster therefore combined lower performance with a stronger impact of irrelevant information.

As a consequence of the psychometric curve shifts (Figure 9), the Point of Subjective Equality (PSE) also varied as a function of irrelevant-stimulus intensity (Figure 10).

**Figure 10:**
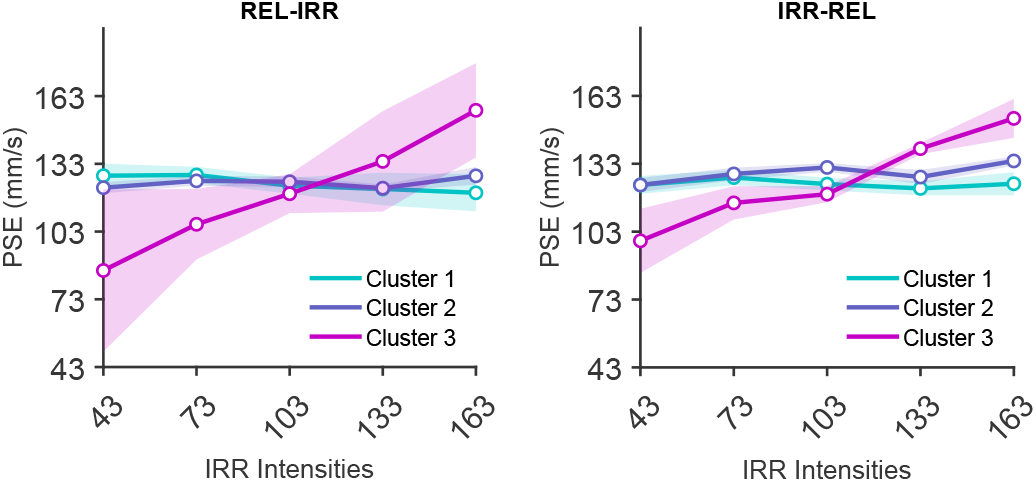
PSE variation as a function of irrelevant-stimulus intensity. PSE values are shown separately for REL-IRR (left) and IRR-REL (right) trials.

Cluster 1 showed a slight decrease in PSE during REL-IRR trials and relatively stable PSE values during IRR-REL trials, consistent with the modest attractive bias observed in the psychometric functions. Cluster 2 exhibited a positive relationship between irrelevant-stimulus intensity and PSE during IRR-REL trials, in agreement with the repulsive behavioral bias identified in this condition. Cluster 3 showed the largest PSE shifts, particularly during REL-IRR trials, mirroring the strong repulsive bias observed in the psychometric data.

The PSE analysis therefore provides a convergent summary of the cluster-specific psychometric shifts.

#### Clusters differ in history dependence and learning dynamics

The cluster-specific psychometric functions also reflected the differences in previous-trial choice dependence captured by the GLM-based clustering (Figure 11).

**Figure 11:**
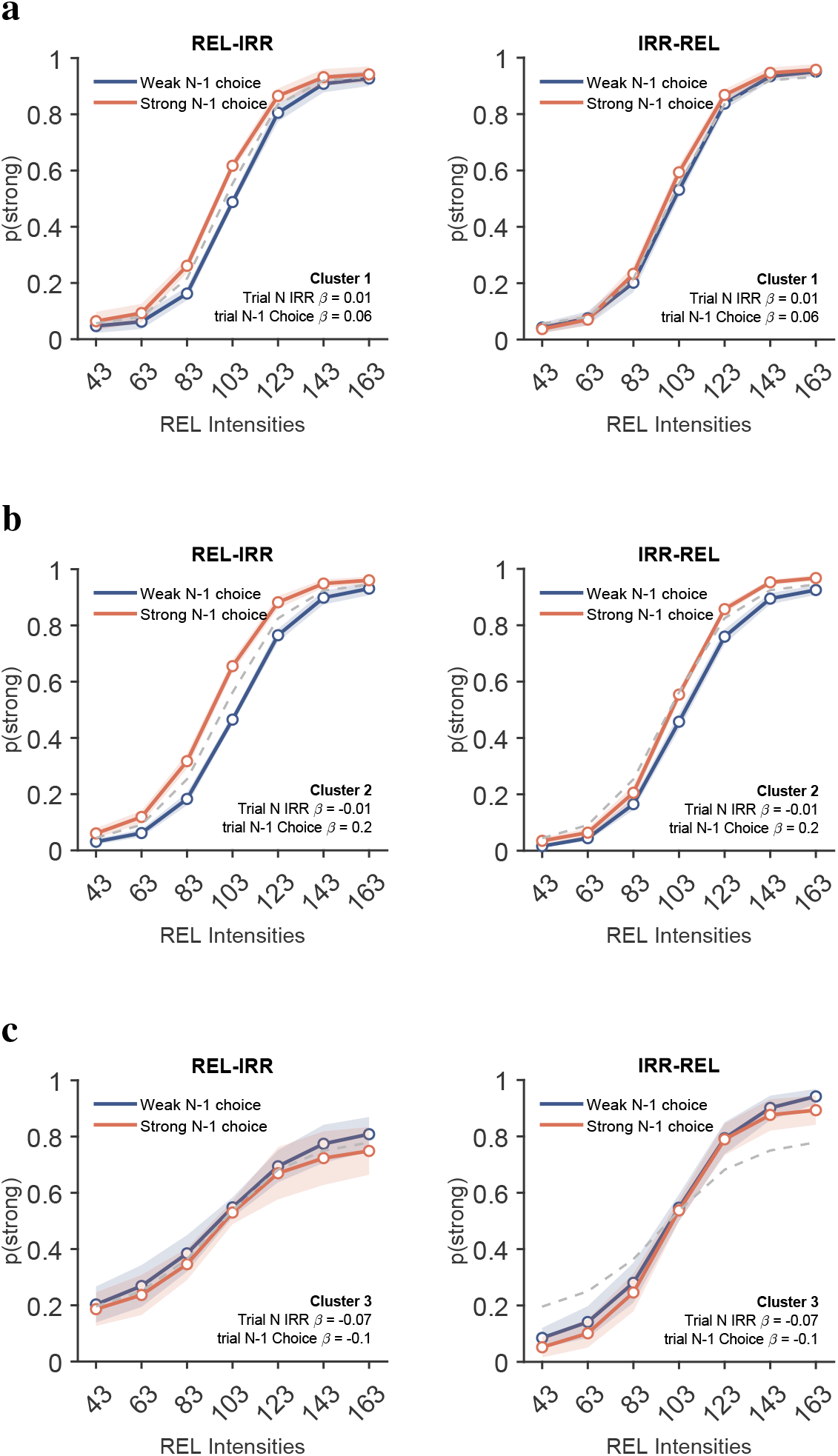
Effect of previous-trial choice at the cluster level. **a.** Psychometric functions conditioned on previous-trial choice for Cluster 1. **b.** Same analysis for Cluster 2. **c.** Same analysis for Cluster 3.

Cluster 2 showed a modest attractive previous-trial choice effect, particularly during IRR-REL trials, consistent with the feature used for clustering. In contrast, Cluster 1 and Cluster 3 showed little evidence of systematic dependence on the immediately preceding choice. Thus, previous-trial history contributed differentially across participant groups and was not uniformly expressed throughout the sample.

Overall, these results show that the apparent homogeneity observed at the population level conceals distinct behavioral profiles. The three clusters exhibited distinct patterns of susceptibility to irrelevant information, ranging from minimal IRR influence in Cluster 1 to increasingly pronounced repulsive effects in Clusters 2 and 3. The third cluster also was characterized by lower task performance. These findings reveal substantial individual differences in how participants filtered irrelevant information and performed perceptual judgments during the dynamic task.

The influence of previous-trial choices differed substantially across the three behavioral clusters. While group-level analyses suggested only a weak effect of trial history on current decisions, cluster-level analyses revealed that this apparent homogeneity concealed marked individual differences.

Cluster 2 exhibited the strongest previous-trial choice effect (Figure 11b). In this group, psychometric functions conditioned on the choice made in trial N-1 showed a modest attractive bias, indicating that participants were more likely to repeat the category selected on the preceding trial. This effect was present in both trial types and was particularly pronounced during REL-IRR trials.

In contrast, Cluster 1 showed little evidence of a systematic influence of previous-trial choice. Psychometric functions corresponding to the two previous-choice conditions largely overlapped, suggesting that current perceptual judgments were determined primarily by information available within the current trial rather than by recent decision history.

Cluster 3 similarly exhibited only weak history dependence. Despite displaying the strongest effects of irrelevant sensory information in other analyses, participants belonging to this cluster showed minimal shifts in psychometric functions as a function of previous-trial choice, indicating that trial history exerted only a limited influence on their decisions.

These results are consistent with the clustering analysis itself, which identified previous-trial choice weight as one of the principal dimensions separating participants. Importantly, the history effects were not distributed uniformly across the population. Instead, they were driven largely by Cluster 2, whereas Clusters 1 and 3 showed little evidence of systematic attraction toward previous choices.

Taken together, these findings demonstrate that history-dependent effects are not a universal feature of task performance. Rather, sensitivity to previous choices characterizes a specific subset of participants, suggesting that the contribution of trial history to perceptual decision-making varies substantially across individuals.

Learning dynamics differed across the three behavioral clusters, indicating that participants not only reached different behavioral states, but also followed different paths to achieve them (Figure 12).

**Figure 12:**
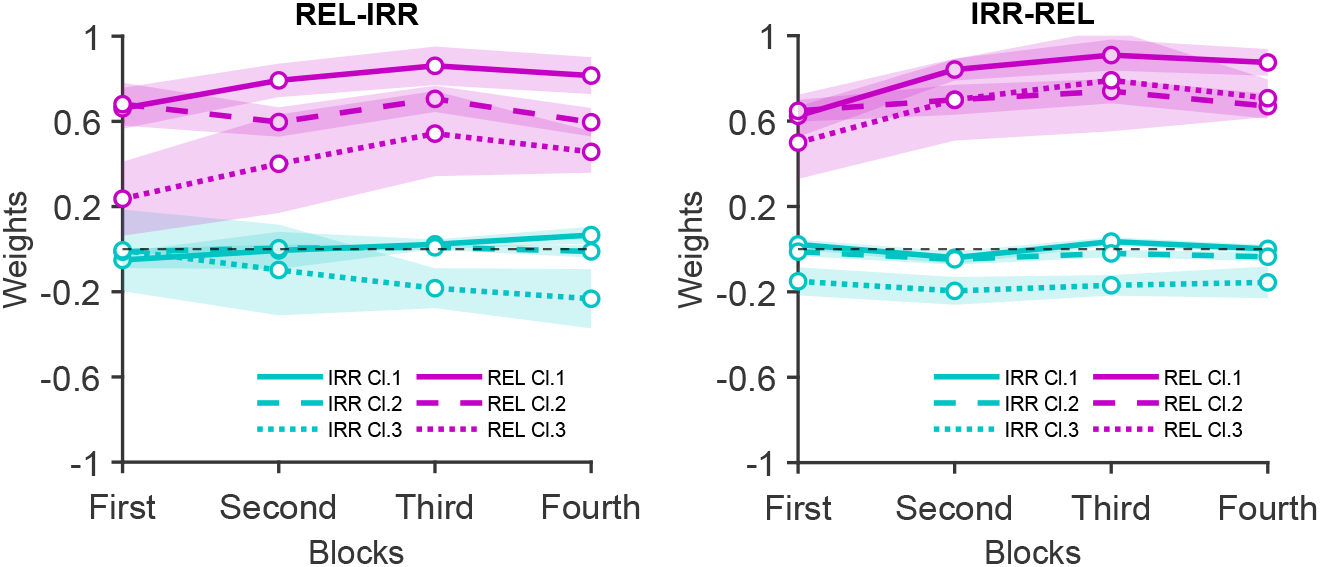
Learning trajectories at the cluster level. GLM weights for relevant (REL) and irrelevant (IRR) stimuli across the four testing blocks, shown separately for REL-IRR and IRR-REL trials.

At the population level, participants showed little evidence of learning because performance was already high at the beginning of training. However, separating participants according to behavioral cluster revealed distinct trajectories in the weighting of relevant and irrelevant information across the four testing blocks.

Cluster 1 showed a gradual increase in the weight assigned to the relevant stimulus in both trial types. In contrast, the weight assigned to the irrelevant stimulus remained close to zero throughout training, particularly during IRR-REL trials. These results indicate a progressive improvement in the use of task-relevant information while maintaining effective suppression of irrelevant inputs.

Cluster 2 exhibited the most stable behavioral profile. Across the four blocks, both relevant- and irrelevant-stimulus weights remained largely unchanged, with irrelevant-stimulus weights remaining close to zero in both trial types. This stability suggests that participants in this cluster adopted an effective strategy early in training and showed little subsequent adjustment in their use of task variables.

Cluster 3 displayed the most distinctive learning trajectory. In IRR-REL trials, the irrelevant-stimulus weights were initially negative, consistent with the repulsive biases observed in the psychometric analyses. Across training, these weights remained relatively stable. In the case of REL-IRR trials, the irrelevant-stimulus weights were initially close to zero, and became increasingly negative over time, indicating a gradual rise in the influence of irrelevant information.

In both trial types, irrelevant-stimulus weights were negative, consistent with the repulsive biases observed in the psychometric analyses. Across training, these weights remained relatively stable during IRR-REL trials, and became more negative during REL-IRR trials, indicating a gradual worsening of performance in the latter trial type. At the same time, the weight assigned to the relevant stimulus increased, particularly during IRR-REL trials. Thus, participants in this cluster showed evidence of increased reliance on task-relevant information, but inconsistent reduction of the influence of irrelevant stimuli.

Despite these differences, all clusters converged toward greater weighting of the relevant stimulus over the course of training. This convergence suggests that participants progressively refined their use of task-relevant information, although the magnitude and temporal profile of learning varied substantially across clusters.

A modest reduction in relevant-stimulus weights was observed in the final training block across all clusters. Because this effect appeared consistently across groups, it may reflect non-specific factors such as fatigue or reduced engagement during the final phase of the experiment.

### Relationship between behavioral performance and introspection

To assess participants’ subjective experience of the task, we administered a brief post-task questionnaire at the end of the second experimental session. The questionnaire evaluated perceived task difficulty, confidence in performance, the influence of stimulus position, and memory of the reference stimulus. We then compared questionnaire responses with objective behavioral measures, including hit rate, previous-trial choice weight, and irrelevant-stimulus weight.

Behavioral performance showed systematic relationships with subjective evaluations of the task (Figure 13). Participants who reported that the task was difficult tended to achieve higher hit rates than participants who described the task as easy. Likewise, participants who reported that trials were more difficult when the relevant stimulus appeared first generally exhibited higher behavioral performance. In contrast, lower performance was associated with uncertainty regarding the intensity of the reference stimulus.

**Figure 13:**
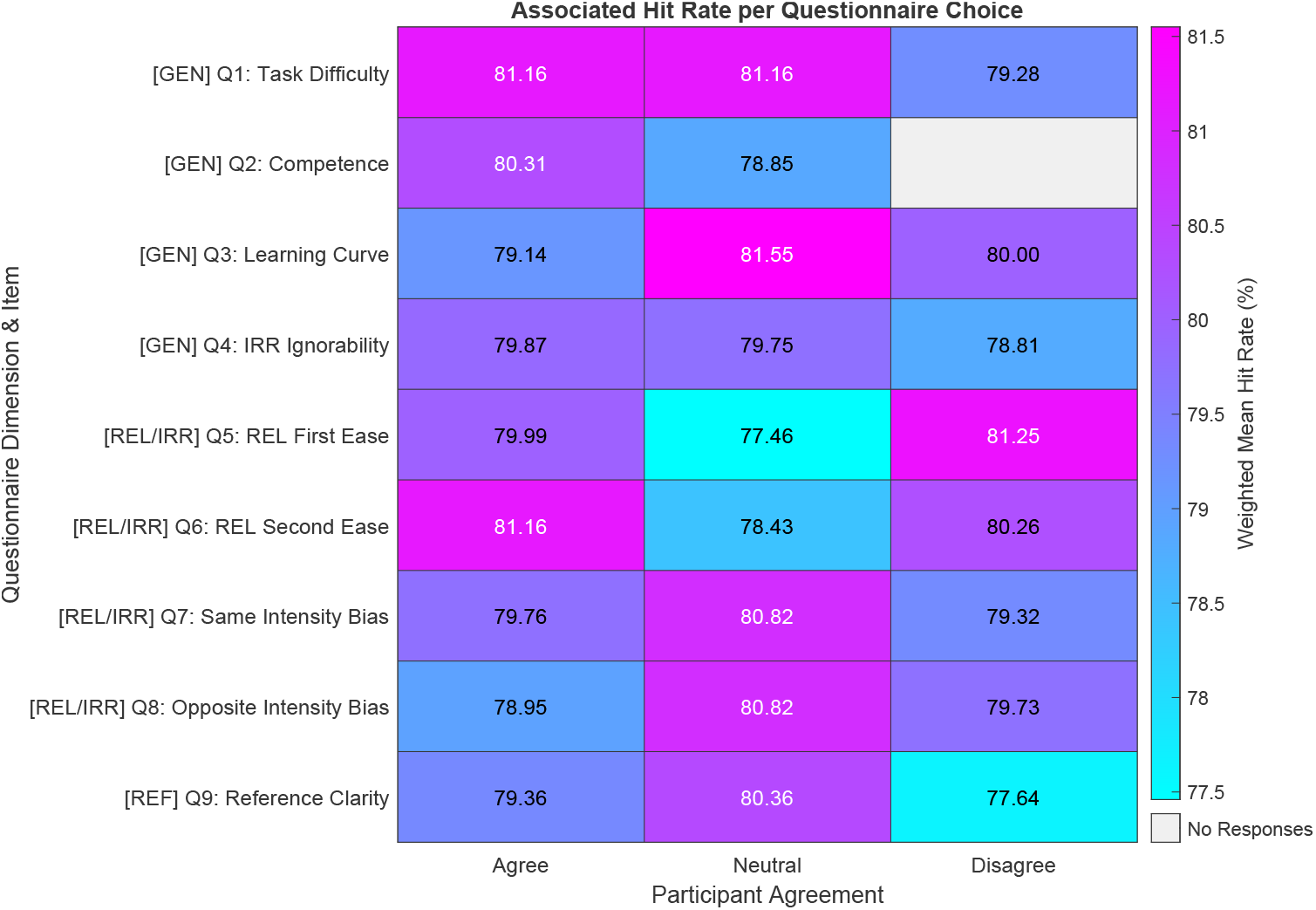
Relationship between task performance and questionnaire responses. Heatmap showing the association between questionnaire responses and average hit rate. Each cell represents a normalized cluster-weighted behavioral measure associated with a specific question-and-answer combination.

A similar pattern emerged when questionnaire responses were examined as a function of previous-trial choice effects (Figure 14). Participants showing stronger previous-trial choice weights were more likely to report that the task was difficult and to identify differences in difficulty between trial types. Conversely, weaker previous-trial choice effects were associated with uncertainty regarding the reference stimulus.

**Figure 14:**
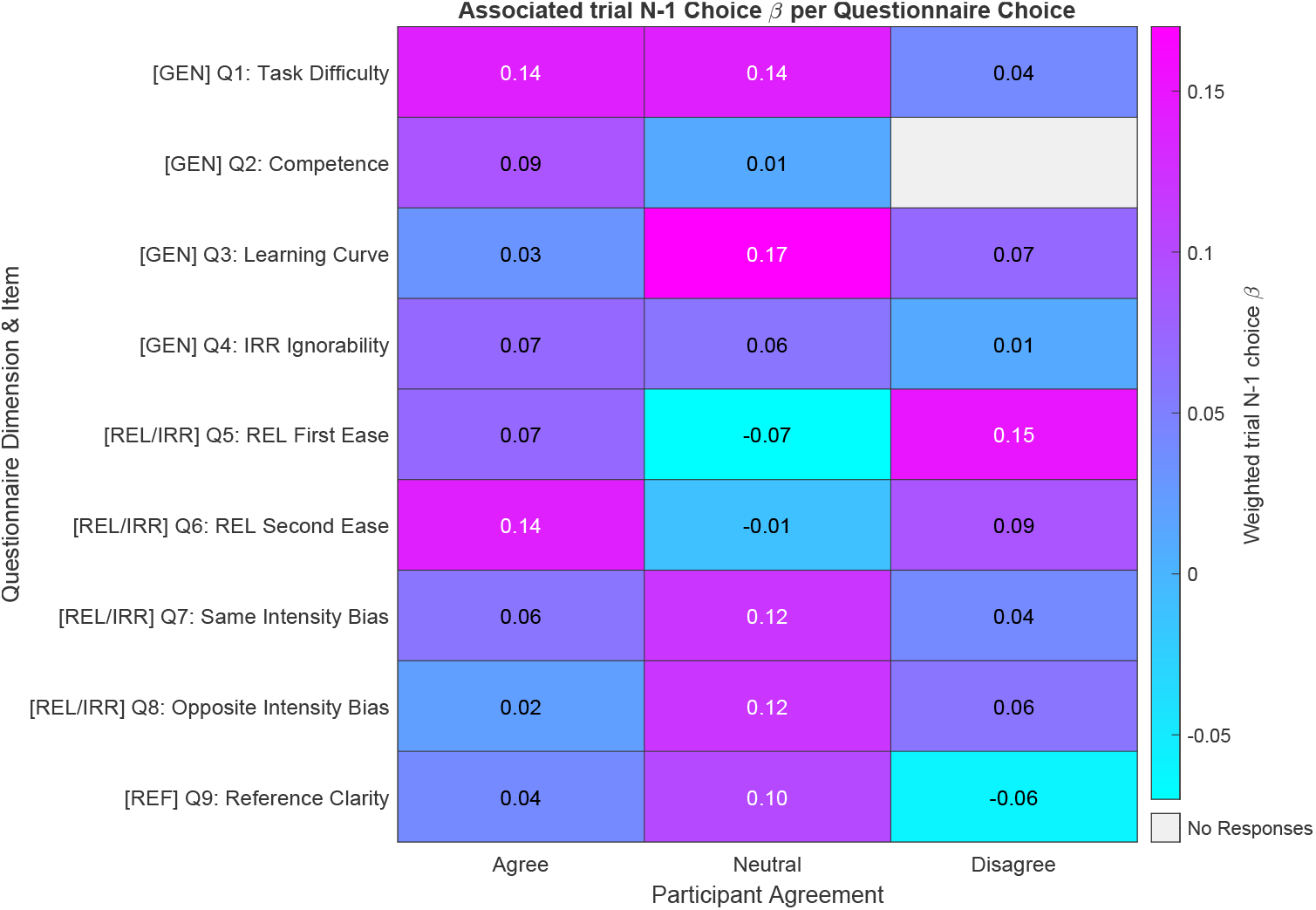
Relationship between previous-trial choice effects and questionnaire responses. Heatmap showing the association between questionnaire responses and average previous-trial choice GLM weight.

Questionnaire responses were also related to the influence of irrelevant information on perceptual judgments (Figure 15). Participants exhibiting smaller irrelevant-stimulus weights generally reported greater task difficulty and were more likely to distinguish between the two trial types. Conversely, participants who reported uncertainty regarding the reference stimulus tended to exhibit larger irrelevant-stimulus effects.

**Figure 15:**
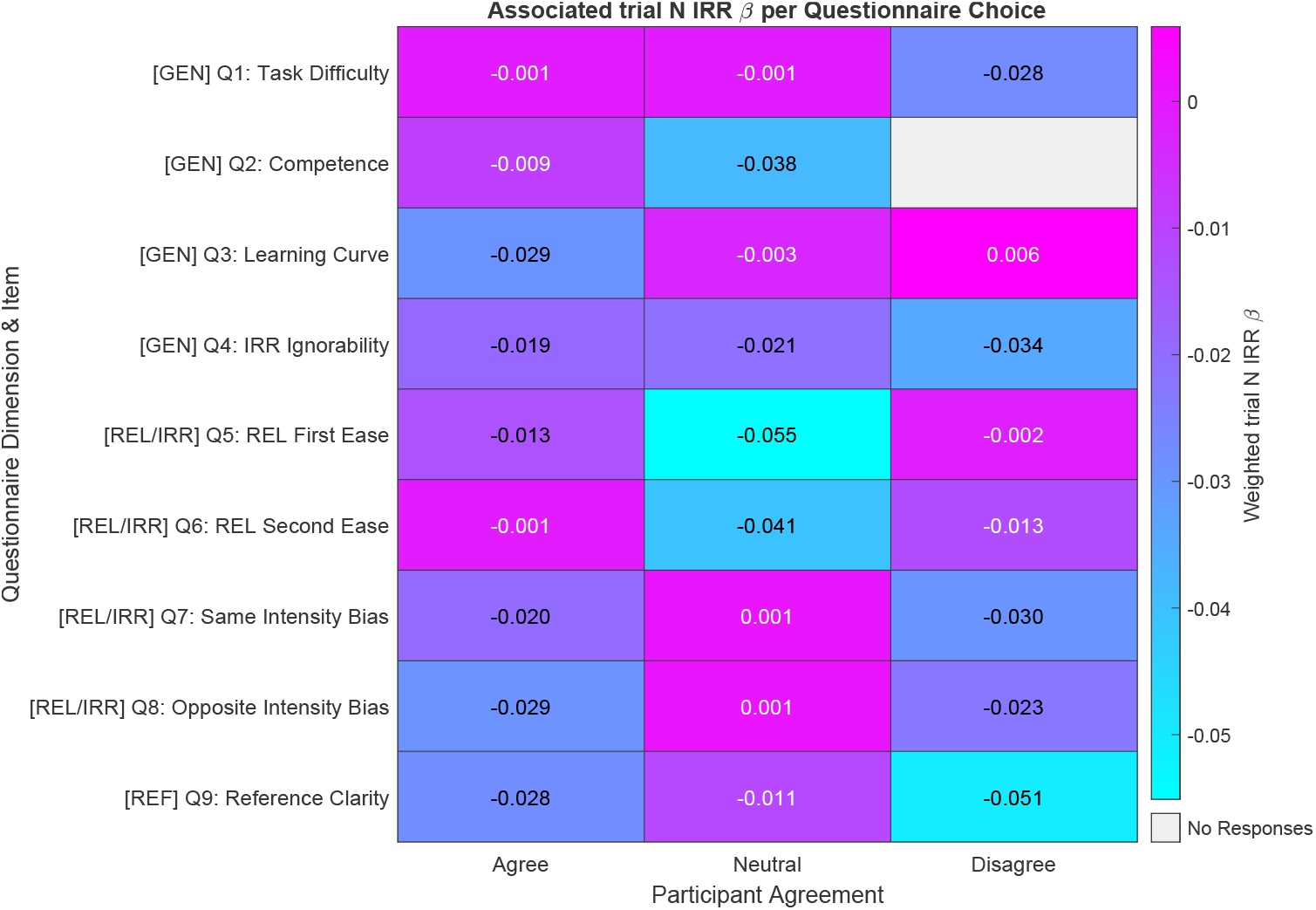
Relationship between irrelevant-stimulus effects and questionnaire responses. Heatmap showing the association between questionnaire responses and average irrelevant-stimulus GLM weight.

Although these analyses are descriptive and exploratory, they reveal a consistent relationship between subjective task awareness and objective behavioral performance. Participants who demonstrated stronger performance, lower susceptibility to irrelevant information, and clearer task representations were also more likely to report greater awareness of task demands and stimulus structure.

Taken together, these findings suggest that individual differences in task performance are reflected not only in behavioral measures but also in participants’ subjective evaluations of the task.

#### Correlation between neuro-divergent traits and attentional control

To investigate whether individual differences in attentional performance were associated with neurodivergent traits, participants completed the AQ-10, SPQ-10, and Aberrant Salience Inventory (ASI) before behavioral testing. We then examined the relationship between questionnaire scores and overall task performance.

No significant correlation was observed between task performance and scores on any of the three questionnaires (Figure 16). This absence of association was evident for measures of autistic traits, sensory-perceptual autistic characteristics, and aberrant salience tendencies.

**Figure 16:**
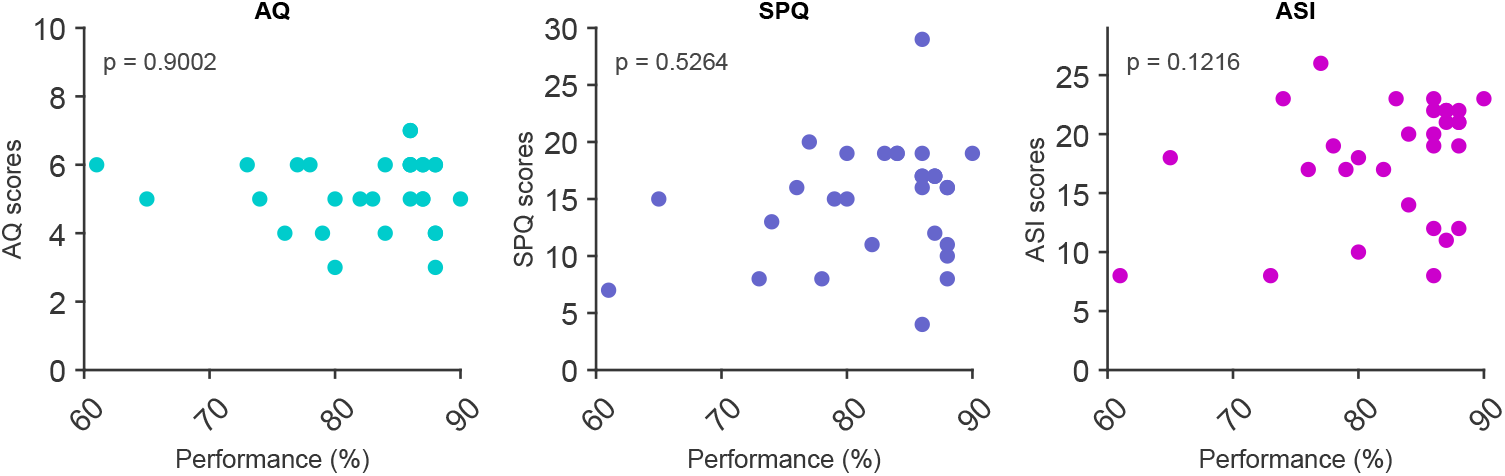
Relationship between questionnaire scores and behavioral performance. Scatterplots showing the relationship between AQ-10 scores (left), SPQ-10 scores (center), ASI scores (right), and average task performance. Each point represents an individual participant. No significant correlations were observed.

These results indicate that variation in questionnaire scores within this sample did not predict success in the dynamic task. Because the present sample consisted exclusively of individuals without reported neuropsychiatric diagnoses, these results should be interpreted as evidence that normal variation in these traits was not associated with meaningful differences in task performance within a non-clinical population.

Overall, the behavioral variability observed throughout the chapter was more strongly explained by differences in task strategy than by variation in questionnaire-derived measures of autistic traits or aberrant salience.

## Discussion

The present study investigated temporal attentional control in the tactile modality of human participants, by asking them to selectively report a relevant tactile stimulus while ignoring an irrelevant tactile distractor presented within the same trial. On average, performance was high. However, analyses conducted at the individual level revealed substantial heterogeneity in behavioral strategies, suggesting that attentional control is not implemented through one single uniform mechanism.

### Individual differences in attentional control

A central finding is that participants could be separated into distinct clusters based on their interaction with irrelevant sensory information. One cluster exhibited slight repulsive biases, where judgments of the relevant stimulus were systematically pulled away from the intensity of the distractor. Another cluster instead showed near-zero influence of irrelevant information. In contrast, a third cluster displayed a strong repulsive influence of the distractor alongside lower overall performance. These results indicate that the same task demands can be solved through qualitatively different behavioral strategies.

These observations parallel our previous findings [14], in which animals exposed to the same task structure exhibited distinct forms of attentional control, characterized by different behavioral and neuronal signatures. The present findings suggest that this variability is not restricted to rodents but may reflect a more general principle governing attentional behavior across species. Rather than representing noise around a common optimal solution, individual differences may correspond to stable alternative strategies for managing competing streams of sensory information.

### Alternative strategies for processing irrelevant information

The residual influence of irrelevant stimuli observed across participants is consistent with an incomplete filtering account of attentional control [21, 22]. Even when explicitly instructed to ignore a stimulus, participants appear unable to eliminate its influence entirely. Instead, information from both tactile events contributes to the final perceptual judgment. Such a mechanism may be advantageous in natural environments, where signals initially considered irrelevant can suddenly become behaviorally meaningful. This scenario suggests that irrelevant stimuli should not be completely suppressed, but should be represented with some salience just below the threshold for action. From this perspective, attentional selection may operate more as a weighting process than as an all-or-none filtering mechanism [23].

The repulsive pattern observed in the third cluster is more difficult to interpret. One possibility is that these participants adopted an active suppression strategy, exaggerating the distinction between relevant and irrelevant information. Such an approach could reduce susceptibility to distraction but may also introduce systematic distortions into the perceptual decision process. Alternatively, the repulsive effect may reflect a less stable learning strategy or a misunderstanding of task contingencies. The fact that this cluster also showed lower average performance favors the possibility that repulsion emerges from a suboptimal solution to the task rather than a superior distractor-suppression mechanism. Nevertheless, because the present study was designed primarily to identify behavioral phenotypes, the cognitive and neural processes underlying these differences remain to be clarified.

The existence of cluster-specific learning trajectories further supports the idea that participants approached the task through distinct computational strategies. Although all individuals experienced the same training procedure, improvements across sessions were not uniform. These differences suggest that attentional strategies emerge dynamically throughout learning rather than being fully established at task onset. Future work employing trial-by-trial computational modeling may help distinguish whether participants differ in sensory encoding, selective weighting of information, decision boundaries, or learning rates.

### Individual traits and metacognitive awareness

One of the motivations of this study was to investigate whether variability in attentional performance could be related to individual differences in autistic traits or aberrant salience [16,24–26]. Contrary to our expectations, neither measure was significantly associated with overall performance nor with the behavioral clusters identified in the task. This absence of effect should be interpreted cautiously. First, the present sample consisted of neurotypical individuals, limiting the range of scores available for detecting meaningful relationships. Second, attentional control is influenced by multiple cognitive factors, and the contribution of broad self-reported traits may be relatively small compared to task-specific strategies. Although null results cannot exclude a relationship between these dimensions and attentional behavior, they suggest that the behavioral heterogeneity observed here is not straightforwardly explained by variation in autistic traits or aberrant salience within a non-clinical population.

We also explored participants’ metacognitive awareness by administering a post-experimental questionnaire assessing subjective evaluations of task difficulty and performance. While these analyses were exploratory, the results suggest that participants possessed varying levels of insight into their own behavior. Individuals reporting greater task difficulty tended to exhibit behavioral patterns consistent with higher objective performance, whereas other participants appeared to overestimate their success. These observations raise the possibility that metacognitive monitoring captures meaningful aspects of attentional strategy selection [27]. Future studies employing established confidence-rating procedures could investigate this relationship more systematically.

## Conclusion

The present study provides important evidence that attentional control in human participants is characterized by substantial individual variability. These findings support the idea that attentional control is not implemented through a single canonical mechanism but rather emerges from multiple behavioral solutions shaped by task demands and individual predispositions. Understanding these alternative strategies may prove essential for developing more comprehensive theories of attention that account not only for average behavior but also for the diversity of ways in which individuals interact with relevant and irrelevant sensory information.

### Limitations and future directions

Several limitations should be acknowledged. First, the sample size may have constrained the detection of subtle associations between questionnaire measures and behavioral performance. Second, the clustering analyses identify reproducible behavioral phenotypes but do not directly explain the underlying cognitive mechanisms. Third, the present work relied exclusively on behavioral measurements and self-report data, preventing direct examination of the neural processes supporting the observed strategies.

## Methods

### Human subjects

30 healthy adult volunteers were recruited to participate in the study. Participants reported normal or corrected-to-normal vision, and no known neurological disorders, as assessed by the demographic questionnaire reporting age, gender, nationality, neurological diagnoses, and current pharmacological treatment. All participants provided informed consent prior to participation. All participants performed two experimental sessions, each lasting approximately one hour and a half, and were paid for their participation, after study completion, on a performance basis. The experimental procedures were conducted in accordance with institutional ethical guidelines and the principles outlined in the Declaration of Helsinki. The study protocol was approved by the Ethics Committee of the International School for Advanced Studies (SISSA). In addition to the behavioral experiment, participants were invited to complete a series of self-report questionnaires assessing autistic traits, aberrant salience, and post-task metacognitive evaluations.

### Experimental apparatus

All experiments took place in the SENSEs Lab, at SISSA. Participants were seated comfortably in front of a computer monitor in a quiet testing environment. Their right arm rested on a pillow placed on a desk. Tactile stimuli were delivered to the fingertip through a vibrotactile stimulator consisting of a plastic probe, attached to a shaker motor (type 4808; Bruel and Kjaer) producing vibrations in the horizontal direction, perpendicular to the fingertip. Visual cues were presented on a computer display positioned directly in front of the participant. Stimulus presentation, response collection, and data storage were controlled using custom LabVIEW software (National Instruments).

### Vibrotactile stimulus generation

Vibrotactile stimuli were vibrations of the probe made of low-pass filtered Gaussian white noise, obtained by stringing together randomly sampled velocities. Velocities were obtained by sampling a normal distribution with 0 mean and defined by the standard deviation *σ*, ranging from 2.84 to 14.85. There were 50 seeds available for each velocity standard deviation. The noise was low-passed through a Butterworth filter with 150 Hz cutoff, amplified, and sent as voltage input to the shaker motor. It is possible to convert the intensities values defined by *σ* in mean speed values (sp, measured in mm/s) by using the following equation:

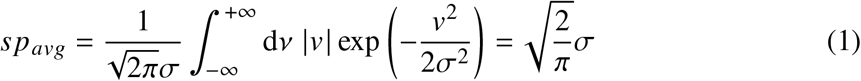

In every session, there were 7 and 5 linearly-spaced values of sp.

### Experimental design

The experiment was based on an adaptation of the Irrelevant Stimulus Paradigm previously developed for rodents [14]. The task design included two types of trials: IRR-REL trials and REL-IRR trials. Delivery of each trial type was randomized within each testing block.

IRR-REL trials were constructed as follows: pre-stimulus delay (500 ms), irrelevant stimulus IRR (500 ms), inter-stimulus interval (1000 ms), relevant stimulus REL (500 ms), post-stimulus delay (500 ms). The visual cue tagging the relevant stimulus lasted 300 ms and overlapped the last 300 ms of the ISI.

REL-IRR trials instead, were designed as follows: pre-stimulus delay (500 ms), relevant stimulus REL (500 ms), inter-stimulus interval (1000 ms), irrelevant stimulus IRR (500 ms), post-stimulus delay (500 ms). The visual cue tagging REL overlapped the last 300 ms of the pre-stimulus delay.

The relevant stimulus set contained 7 intensities, whereas the irrelevant stimulus set comprised 5 intensities. These adaptations from the rodent version provided a compromise between sufficient trial count and total study duration.

The intensity of both relevant and irrelevant stimuli varied across trials according to predefined stimulus sets. Participants responded after the presentation of both tactile stimuli by selecting the category corresponding to the perceived intensity of the relevant stimulus.

### Experimental procedure

The experiment consisted of four test blocks distributed across two experimental sessions. Sessions were separated in time to allow the assessment of learning trajectories and performance changes across training.

Prior to the start of the experiment, participants received verbal and written instructions explaining the task structure. Practice trials were administered to ensure that participants understood the response requirements and the meaning of the visual cues.

During the test blocks, participants completed multiple trials in which relevant stimulus position, stimulus intensity, and distractor intensity were systematically varied.

### Behavioral analysis

#### Psychometric curve estimation

All analyses were performed in MATLAB (MathWorks, Natick MA) using custom code. To estimate psychometric functions, we computed for each stimulus vibration *sp*, the proportion of trials in which the participant responded by pressing the button corresponding to the "strong" category. We fitted the data with a cumulative Gaussian logistic function, including asymmetric lapse parameters:

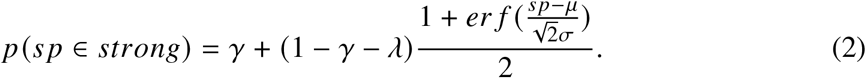

where *sp* is the stimulus intensity, *μ* is the midpoint parameter, *σ* is the slope parameter, *γ* and *λ* the lower and higher asymptotes of the function, respectively, corresponding to lapse rates. Parameters values were estimated by MLE using the MATLAB functions fmincon and fminsearch.

Psychometric functions were fitted separately for each participant. For across-subject plots, fitted probabilities at each tested stimulus intensity were averaged across participants, and S.E.M. was computed across the individually fitted probabilities. Behavioral performance was quantified as the proportion of correct responses across trials. Psychometric analyses were performed to evaluate the influence of stimulus intensity and distractor magnitude on perceptual judgments.

#### Generalized Linear Model estimation

GLM analyses were implemented using custom-made MATLAB code, fitting free parameters with the following equation:

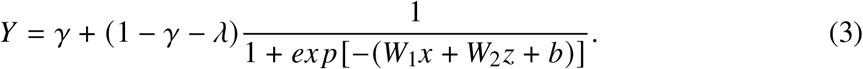

Where *Y* is the predicted participant’s choice, *γ* and *λ* are, respectively, the lower and upper lapse rates, *W*_1_ is the weight for the REL stimulus, *W*_2_ is the weight for the IRR stimulus, *x* is the REL stimulus intensity, and *z* is the IRR stimulus intensity. Parameters values were optimized by MLE using the MATLAB functions fmincon and fminsearch.

For the learning trajectory analyses, GLM weights were estimated for individual participants and individual blocks, and then averaged across subjects.

#### Point of Subjective Equality estimation

We estimated the Point of Subjective Equality (PSE) by computing the psychometric curves and storing the parameter *μ*. PSE was defined as the parameter *μ* of the fitted cumulative Gaussian psychometric function, and expressed in the corresponding stimulus velocity.

#### Clustering analysis

To investigate individual differences in task performance, participants were clustered according to two behavioral features quantifying the influence on current perceptual judgments: the effect of the irrelevant stimulus presented in trial *N* and the effect of the choice made in trial *N* − 1. For each participant, the two clustering features were estimated using separate binomial Generalized Linear Models with a probit link, implemented with the MATLAB fitglm function. The trial *N* irrelevant-stimulus weight was obtained by modeling current-trial choice as a function of relevant- and irrelevant-stimulus intensities, whereas previous-choice dependence was estimated by modeling current-trial choice as a function of the choice made on trial *N* − 1. The corresponding GLM coefficients were retained as the trial *N* IRR and trial *N* − 1 choice features, respectively.

Each participant was therefore represented by a two-dimensional feature vector containing the trial *N* IRR weight and the trial *N* − 1 choice weight. The resulting 30 × 2 feature matrix was clustered using the *k-medoids* algorithm with Manhattan distance (also known as city-block distance). We compared solutions with *k* = 2, *k* = 3, and *k* = 4. The three-cluster solution was retained as a parsimonious separation into qualitatively distinct GLM-weight profiles, whereas increasing *k* to 4 primarily subdivided one of these profiles according to weight magnitude without revealing an additional qualitatively distinct pattern.

The *k*-medoids clustering minimizes the within-cluster distance between each participant and the medoid of the cluster to which that participant is assigned:

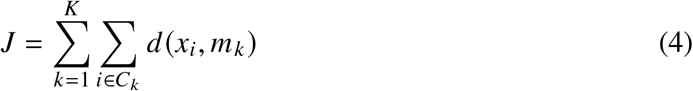

where *K* is the total number of clusters (*K* = 3), *C_k_* is the set of participants assigned to cluster *k*, *x_i_* is the two-dimensional feature vector describing participant *i*, and *m_k_* is the medoid of cluster *k*, corresponding to an actual participant in the dataset.

Manhattan distance was used as the distance metric:

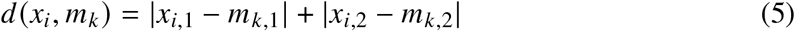

where *x_i_*_,1_ and *m_k_*_,1_ correspond to the trial *N* IRR GLM weights of participant *i* and the cluster medoid, respectively, whereas *x_i_*_,2_ and *m_k_*_,2_ correspond to their trial *N* − 1 choice GLM weights.

Cluster assignments were subsequently used to characterize performance, learning trajectories, and cluster-specific psychometric patterns

### Questionnaire Measures

#### Autism Spectrum Quotient and Sensory-Perception Quotient

Participants completed the AQ-10 to assess autistic traits and the SPQ-10 to assess sensory-perceptual characteristics associated with autism ( [28]). Total scores were calculated according to the corresponding scoring procedures and used as continuous measures of trait variation within the sample. Total questionnaire scores were correlated to the behavioral performance (i.e., proportion of hit trials) using Spearman’s correlation, with *α* = 0.05.

#### Aberrant Salience

Participants completed a questionnaire assessing aberrant salience tendencies, defined as the propensity to attribute significance to otherwise neutral events or stimuli (ASI questionnaire, [29]). Total questionnaire scores were correlated to the behavioral performance (i.e., proportion of hit trials) using Spearman’s correlation, with *α* = 0.05.

#### Metacognitive Assessment

Following completion of the experimental sessions, participants completed a brief questionnaire evaluating their subjective experience of the task. Items assessed perceived task difficulty, confidence in performance, and general impressions regarding attentional demands.

These measures were analyzed exploratively to examine possible relationships between self-evaluation and objective task performance. The procedure involved computing a weighted average of the behavioral metrics across a cluster-based feature space.

First, the distribution of question-and-answer responses across the three clusters was tabulated: to prevent cluster size imbalances from biasing the results, these raw counts were normalized by total cluster size to yield a within-cluster response proportion. Next, these normalized proportions were used as weights to calculate a weighted average of the three behavioral metrics (hit rate, trial N IRR *β*, trial N-1 choice *β*) for each individual cell.

Each cell in the resulting heatmaps reflects the normalized, cluster-weighted behavioral metric associated to a specific item and answer of the questionnaire. Below, the equation describing the measure contained in each cell:

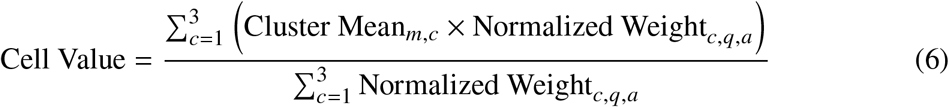

Where *c* represents the cluster index (*c* ∈ {1, 2, 3}). *m* represents the specific behavioral or computational metric evaluated (Hit Rate, trial *N* IRR *β*, or trial *N* −1 choice *β*). *q* represents the specific item (question) from the questionnaire (*q* ∈ {1,…, 9}). *a* represents the participant’s level of agreement (answer) for that item (*a* ∈ {Agree, Neutral, Disagree}). Cluster Mean*_m_*_,*c*_ is the mean value of metric *m* within cluster *c*. Normalized Weight*_c_*_,*q*,*a*_ is the response proportion of cluster *c* associated with item *q* and answer *a*, computed as the raw count of participants in cluster *c* who chose answer *a* divided by the total number of individuals assigned to that cluster.

## Author contribution

M.G. and M.E.D. designed the experiment. M.G. devised the data collection protocol, trained, and supervised the data collector. M.G. analyzed the data. M.G. and M.E.D. interpreted and discussed the results. M.G. and M.E.D. wrote the paper. M.E.D. performed the acquisition of fundings.

## Acknowledgments

We acknowledge the financial support of the the Human Frontier Science Program (http://www.hfsp.org; project RGP0017/2021), the financial support of the Italian Ministry PRIN 2022 funding (contract 20224FWF2J), the financial support of the Italian Ministry PNRR funding (contract P20229752W), the financial support of the Regional Laboratory for Advanced Mechatronics, LAMA FVG (http://lamafvg.it), and the financial support of the European Union Horizon 2020 MSCA Programme (grant 813713). We thank Yoanna Fernandez for collecting the behavioral data. We also thank Fabrizio Manzino (CyNexo) for the invaluable technical assistance for and for software development.

## Supplementary Material

### Psychometric functions

The psychometric functions reported in the main figures were obtained by averaging the fitted probabilities estimated separately for each participant. The following supplementary figures instead show the corresponding empirical probabilities, without fitting. Shaded areas represent the S.E.M. across individually computed empirical probabilities. The lines connect consecutive empirical probabilities for visualization.

**Figure S1:**
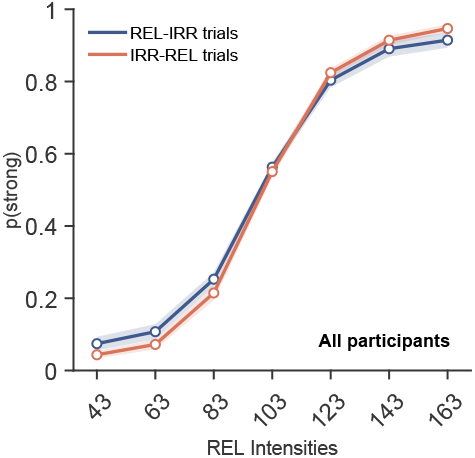
Across-subject overall performance during the dynamic task. Psychometric functions showing average performance across participants for REL-IRR and IRR-REL trials.

**Figure S2:**
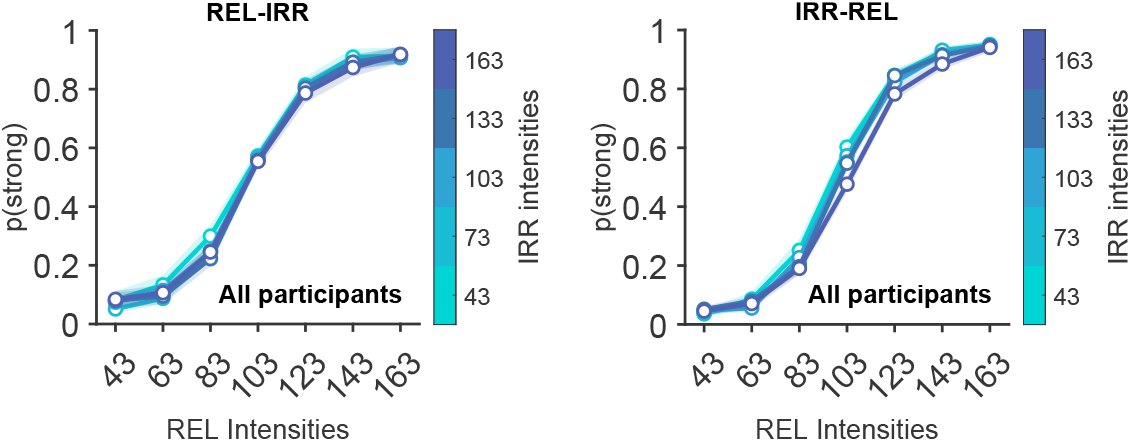
Across-subject irrelevant-stimulus bias during the dynamic task. Psychometric functions conditioned on irrelevant-stimulus intensity during REL-IRR trials (left) and IRR-REL trials (right).

**Figure S3:**
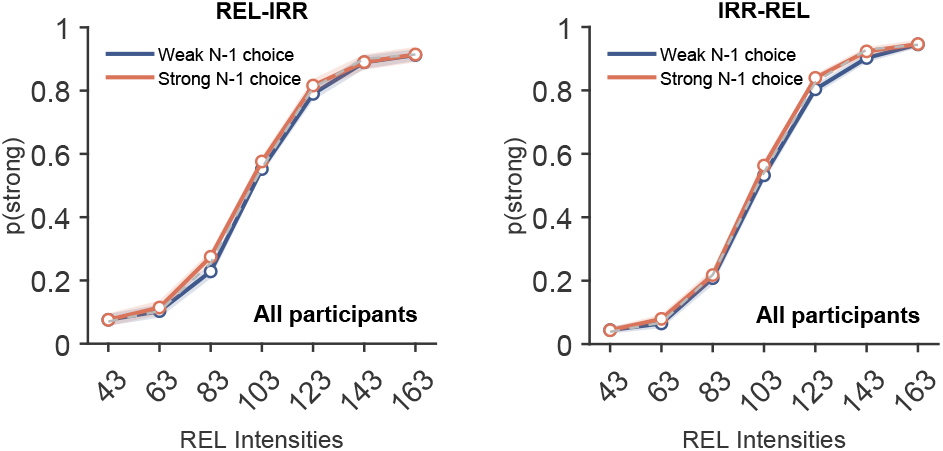
Effect of previous-trial choice on current perceptual judgments. Psychometric functions conditioned on the choice made in trial N-1, shown separately for REL-IRR trials (left) and IRR-REL trials (right).

**Figure S4:**
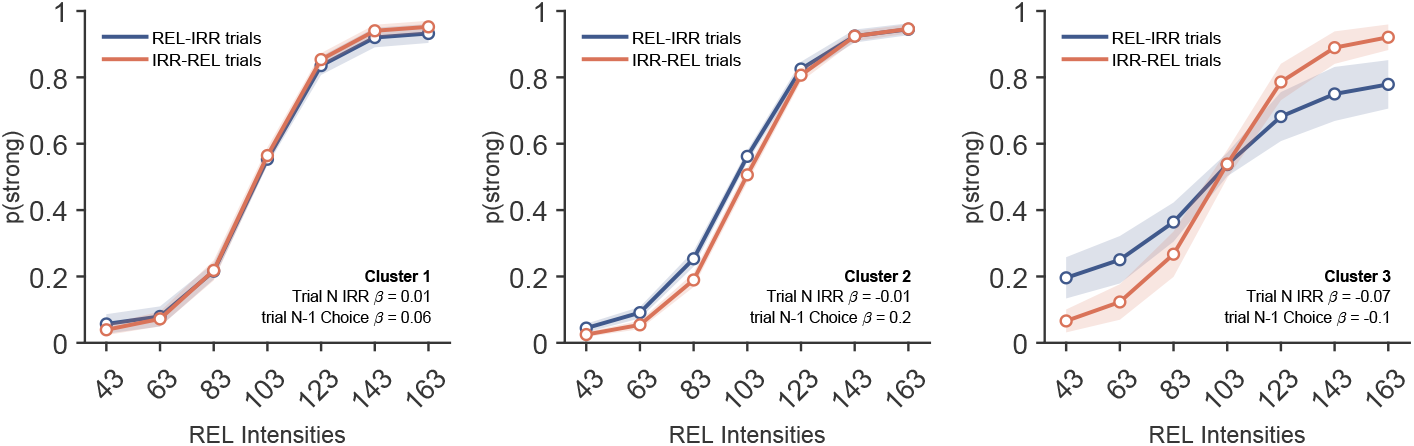
Average performance at the cluster level. Psychometric functions showing average task performance for Clusters 1, 2, and 3.

**Figure S5:**
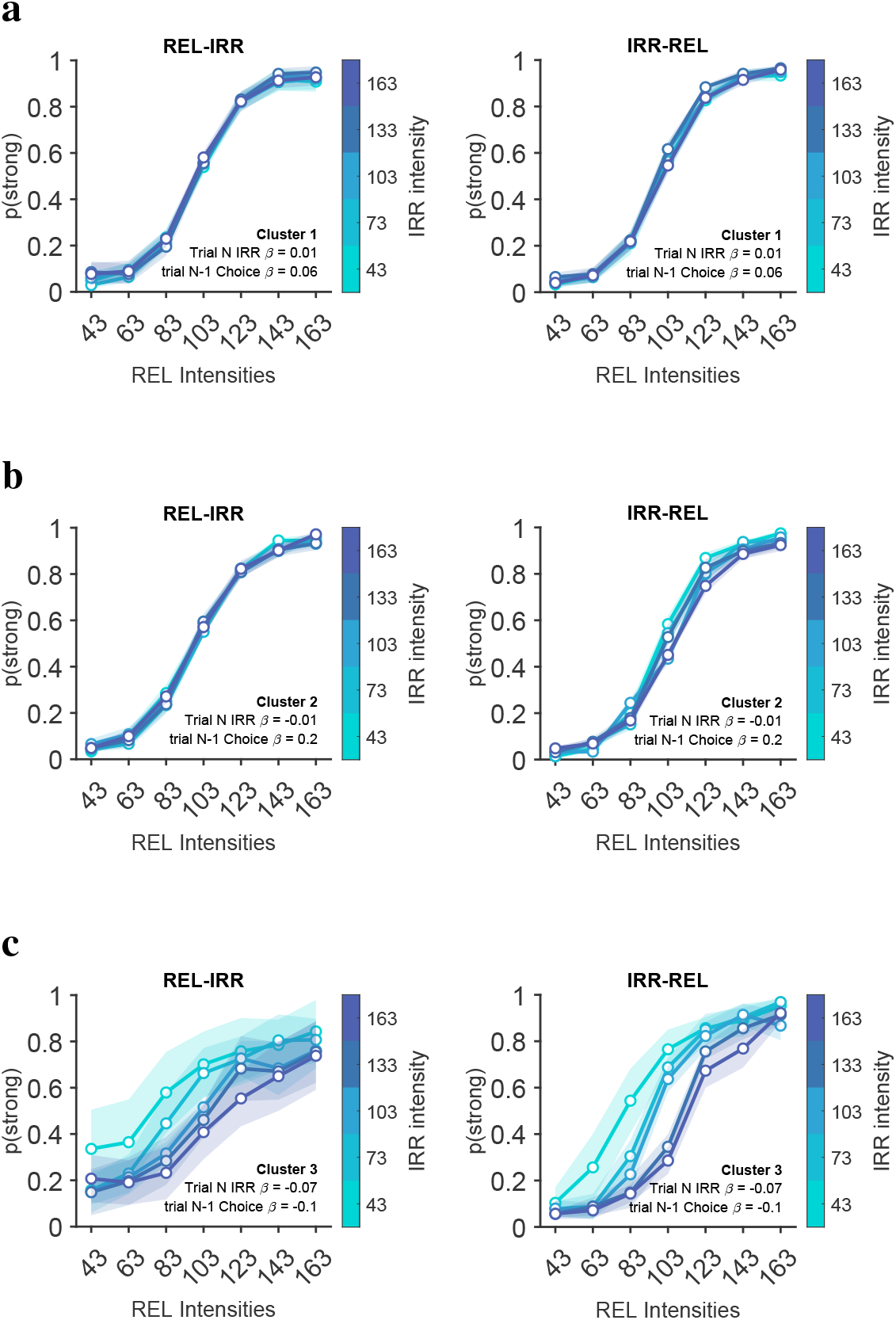
rrelevant-stimulus bias at the cluster level. **a.** Psychometric functions conditioned on irrelevant-stimulus intensity for Cluster 1.**b.** Same analysis for Cluster 2. **c.** Same analysis for Cluster 3.

**Figure S6:**
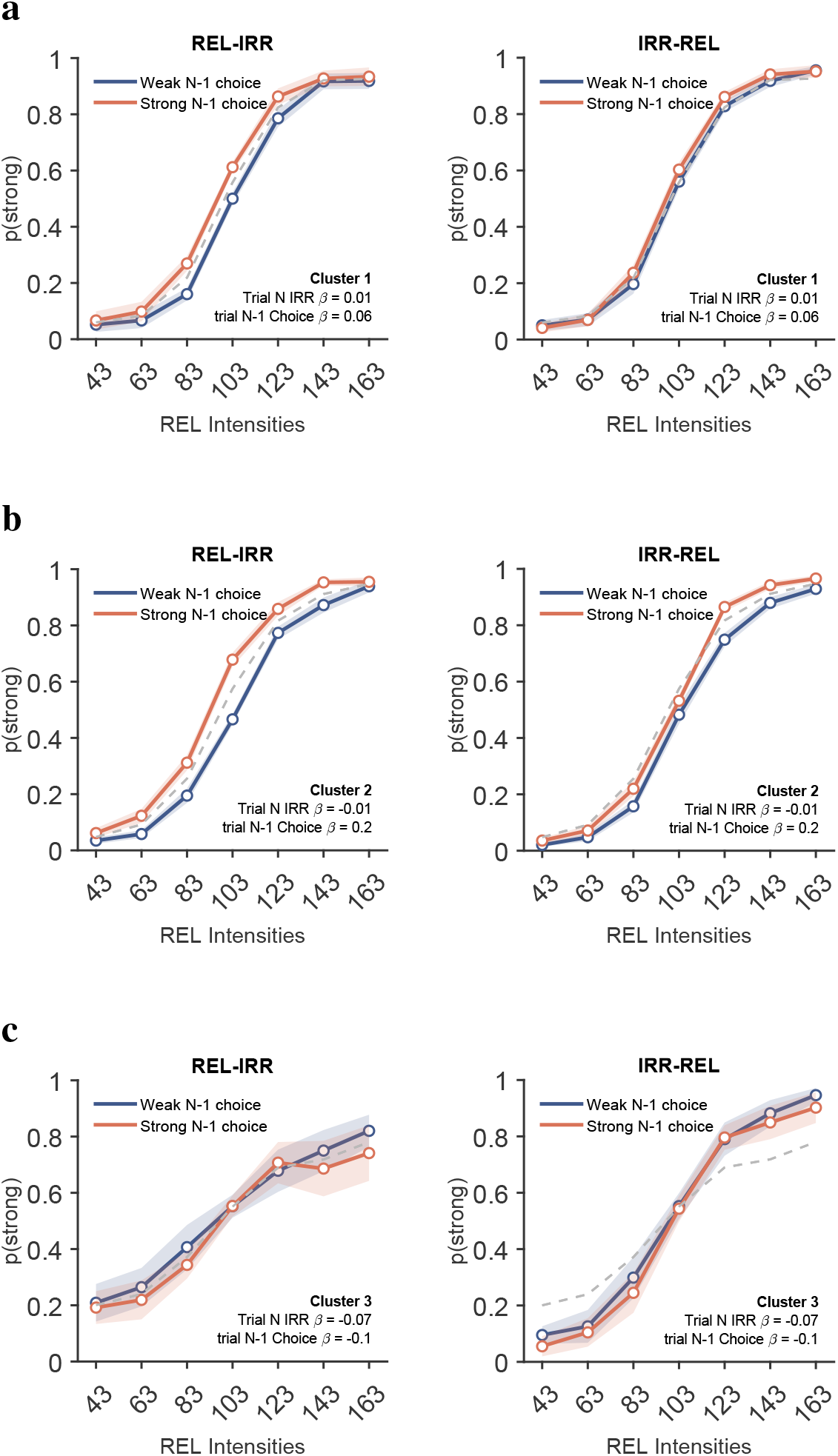
Effect of previous-trial choice at the cluster level. **a.** Psychometric functions conditioned on previous-trial choice for Cluster 1. **b.** Same analysis for Cluster 2. **c.** Same analysis for Cluster 3.

### Final questionnaire

**Figure S7:**
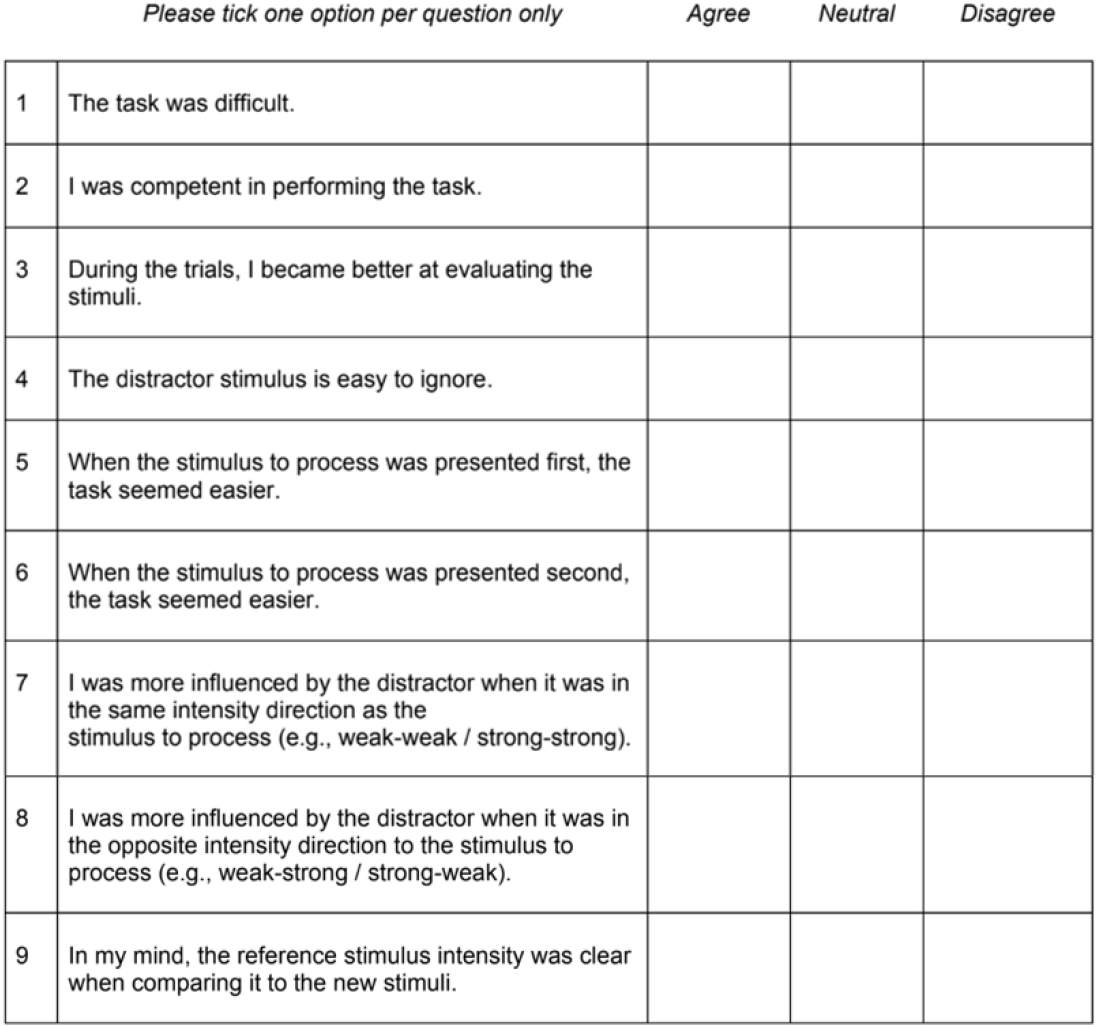
Final questionnaire administered to human participants. The participants received the Italian version of the questionnaire, since they all spoke Italian as mother tongue.

## Notes

### Competing Interest Statement

The authors have declared no competing interest.

